# Modulating social learning-induced evaluation updating during human sleep

**DOI:** 10.1101/2023.12.11.571114

**Authors:** Danni Chen, Tao Xia, Ziqing Yao, Lingqi Zhang, Xiaoqing Hu

## Abstract

People often change their evaluations upon learning about their peers’ evaluations, i.e., social learning. Given sleep’s vital role in consolidating daytime experiences, sleep may facilitate social learning and thereby further changing people’s evaluations. Combining a social learning task and the sleep-based targeted memory reactivation technique, we asked whether social learning-induced evaluation changes can be modulated during sleep. After participants indicated their initial evaluation for snacks, they learned about their peers’ evaluation while hearing the snacks’ spoken names. During the post-learning non-rapid-eye-movement sleep, we re-played half of the snack names (i.e., cued snack) to reactivate the associated peers’ evaluations. Upon waking up, we found that the social learning-induced evaluation changes further enlarged for both cued and uncued snacks. Examining sleep electroencephalogram (EEG) activity revealed that cue-elicited delta-theta EEG power and the overnight N2 sleep spindle density predicted post-sleep evaluation changes for cued but not for uncued snacks. Our findings suggested that sleep-mediated memory reactivation processes could strengthen social learning-induced evaluation changes.

## Introduction

Evaluations and choices are often guided by retrieval of first-hand experiences: when choosing a restaurant, we often think about our last visit, the dining experiences, and the accompanying emotional feelings (Amodio, 2019; Biderman et al., 2020; Hütter, 2022). However, in addition to using first-hand experiences to guide our choices (Murty et al., 2016; Wimmer & Büchel, 2016; Wimmer & Shohamy, 2012), we also acquire or change evaluations via observing our peers’ evaluations and choices, known as social learning (Berns et al., 2010; Campbell-Meiklejohn et al., 2010; Kendal et al., 2018). Social learning is prevalent in society, influencing everyday choices, such as purchasing snacks or books, and even sacred moral values (Brady et al., 2021; Nook & Zaki, 2015; Yu et al., 2021; Zaki et al., 2011). Specifically, social learning can be induced in lab settings: following observing peers’ evaluations, participants often change their initial evaluations (Chen et al., 2023; Huang et al., 2014; Nook & Zaki, 2015; Zaki et al., 2011). These social learning-induced evaluation changes can even last for days after the learning (Huang et al., 2014; Izuma & Adolphs, 2013). The observed long-term effect raises an intriguing yet untested question: how does memory consolidation during post-learning sleep influence the social learning effect?

Mounting evidence suggests that sleep consolidates recently acquired memories via covert memory reactivation processes (Brodt et al., 2023; Klinzing et al., 2019; Rasch & Born, 2013). Employing a method known as Targeted Memory Reactivation (TMR), researchers can initiate and guide covert memory reactivation during sleep to promote memory consolidation (Oudiette & Paller, 2013; Paller et al., 2021). This TMR procedure typically consists of three phases: 1) pre-sleep learning, participants would learn materials accompanying sensory cues (e.g., auditory tones, spoken words, odor); 2) TMR during sleep, during which the experimenter will re-present the same sensory cues (i.e., memory reminders) to sleeping participants to reactivate the associated memories; and 3) post-sleep tests, upon awakening, participants would complete tests to assess the impact of TMR. Accumulating evidence has demonstrated that TMR benefits various types of memories (for a meta-analysis, see Hu et al., 2020), including speech-word pair associative learning (Cairney et al., 2017), skills learning (Antony et al., 2012; Rakowska et al., 2021), spatial memories (Rudoy et al., 2009; Shanahan et al., 2018), and emotional memories (Lehmann et al., 2016; Yuksel et al., 2023). Here, we aimed to explore the potential impact of TMR on people’s evaluations acquired through prior social learning.

To date, only a few studies have explored the potential impact of sleep and/or TMR on evaluation. For example, sleep (vs. wakefulness) promoted adaptive evaluative choices, by strengthening evaluative learning memories (Jin et al., 2023). Employing TMR, research shows that re-playing snacks’ spoken names during non-rapid eye movement (NREM) sleep could augment subjective preferences for these snacks (Ai et al., 2018). Moreover, re-playing the sound cues paired with the prior counter-bias training during NREM sleep further reduced implicit social biases (Hu et al., 2015; but see Humiston & Wamsley, 2019). These findings suggest that sleep and/or TMR could modulate evaluations and choices, potentially through sleep-mediated reactivation of pre-sleep evaluative learning memories.

Analyzing cue-elicited electroencephalogram (EEG) activity during sleep can provide insights into the underlying neural mechanisms of TMR. Specifically, cue-elicited delta (1-4 Hz) and theta (4-8 Hz) activities have been shown to predict TMR benefits on memory performance (Liu et al., 2023; Oudiette et al., 2013; Rihm et al., 2014; Schreiner et al., 2015; Xia, Chen, et al., 2023). Notably, previous research also revealed the role of cue-elicited delta and theta power in predicting TMR benefits in evaluation updating (Ai et al., 2018; Xia, Antony, et al., 2023). Furthermore, substantial evidence has indicated that overnight sleep spindle is implicated in memory re-processing during sleep (Antony et al., 2019; Clemens et al., 2005; Kurdziel et al., 2013; Mednick et al., 2013) and predicts the TMR benefits (Creery et al., 2015; Xia, Antony, et al., 2023). We thus investigated the neural mechanisms, focusing on the delta/theta power and the sleep spindles underlying the reactivation of daytime social learning experiences.

In the present study, we employed the TMR to investigate how reactivating prior social learning experiences during NREM sleep would influence subsequent evaluation. Following the initial evaluation for snacks, participants learned their peers’ evaluations as feedback while listening to the snacks’ spoken names. Via multiple learning rounds, these spoken names would serve as memory reminders about peers’ evaluations of the snacks. During the subsequent NREM sleep, we re-played half of these spoken names to reactivate the associated peers’ evaluations. Upon waking up, participants showed enlarged social learning-induced evaluation updating for both cued and uncued snacks. Accompanying behavioral changes, cue-elicited delta-theta EEG power, and the overnight N2 spindle density were associated with the evaluation updating for cued but not for uncued snacks. These results suggested that sleep-mediated memory consolidation processes may fortify social learning-induced evaluation updating.

## Methods

### Participants

We recruited 45 participants from a local university (35 females; Age, *Mean* = 22.98, *S.D.* = 2.81). Participants were excluded from subsequent behavioral and EEG analysis if the auditory cues were played fewer than four rounds (*n* = 9) or due to technical problems during EEG recording (*n* = 2), resulting in 34 participants being included in the analyses. All participants were native Chinese speakers, right-handed, not color-blind, and had normal or correct-or-normal vision. In addition, they reported good sleep qualities without any history of neurological, psychiatric, or sleep disorders. All participants provided written informed consent prior to the participation and were debriefed and compensated after they completed the study. This research was approved by the Human Research Ethics Committee of the University of Hong Kong (HREC No. EA1904004).

### Stimuli

We selected 48 snack images from the snack and food images database (Hare et al., 2011; Plassmann et al., 2007). Spoken names of snacks were generated in English using the Microsoft Azure Text-to-Speech function (language = “en-US”). The 48 snacks were then allocated to one of six experimental conditions based on each participant’s baseline evaluation (i.e., the preference rating before the social learning). To do this, all 48 snacks were first sorted in descending order based on the baseline ratings and were subsequently divided into eight subgroups following this ranked order, each consisting of six snacks. For instance, snacks in this first subgroup would rank from first to sixth, while snacks in the second subgroup would rank from seventh to twelfth, and so on. Next, the six snacks in each subgroup were randomly assigned to one of the six experimental conditions in 2 (TMR: cued vs. uncued) by 3 (Peer’s evaluation feedback in social learning task: lower vs. consistent vs. higher) design. This procedure resulted in eight items in each of the six experimental conditions, with baseline preferences and familiarity ratings not significantly different between different conditions (*p*s > .087; see Table S1 for details).

### Design and Procedure

#### Procedure

All tasks were programmed and presented by PsychoPy (2020.1.3) (Peirce et al., 2019). Participants visited the lab twice, separated by three days (Figure 1A).

**Figure 1:**
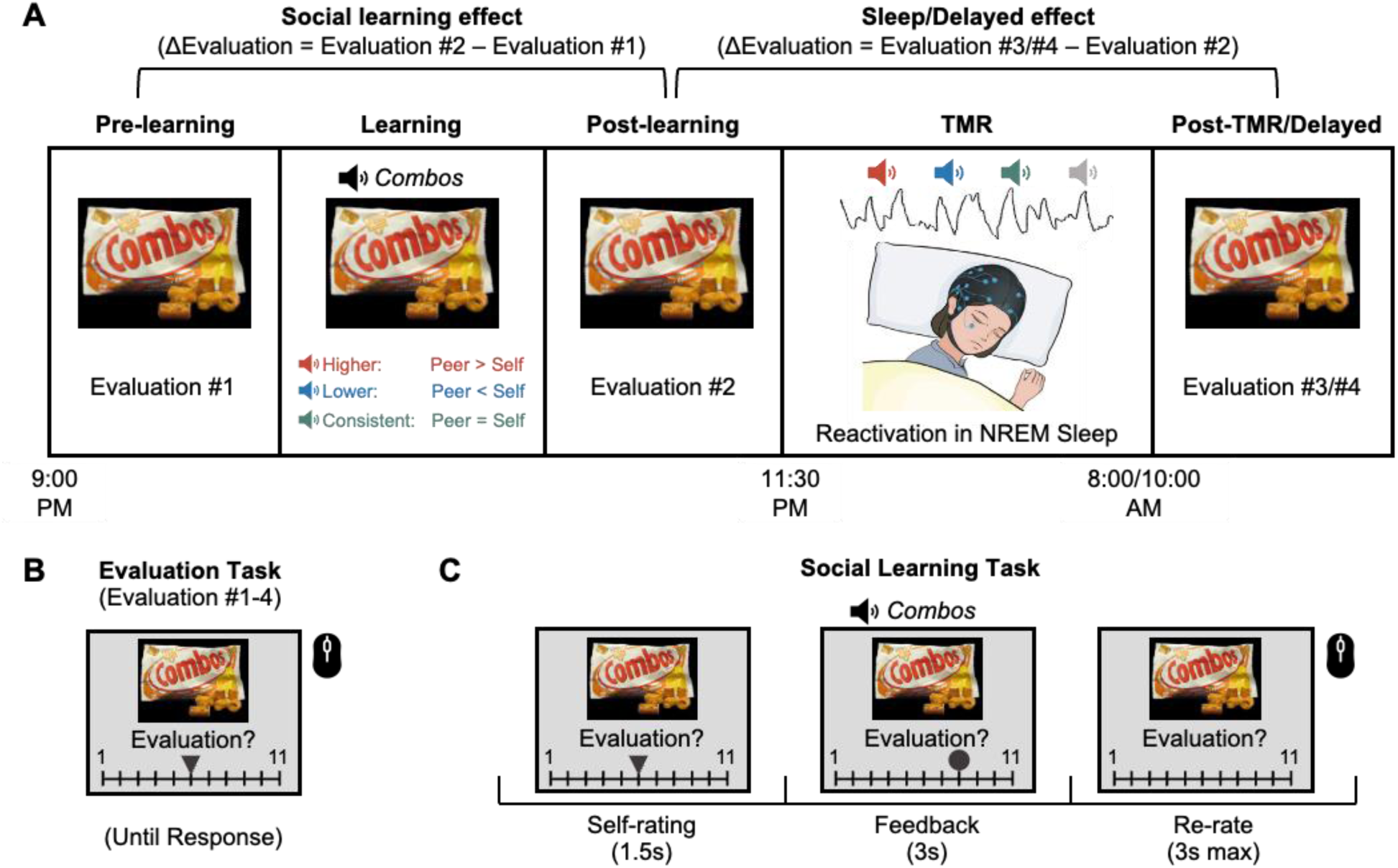
A flowchart of the experiment procedure. A) The experiment included pre-learning baseline tests, social learning of peers’ evaluation task, post-learning tests, TMR during NREM sleep, post-TMR immediate and 3-day delayed tests. B) An exemplar trial in the Evaluation tasks: Participants evaluated each of the 48 snacks using a mouse clicking on a 1-11 scale, ranging from not preferred at all to most preferred.. C) During the Social Learning task, participants learned the evaluation from their peers (a circle indicating their peers’ evaluation). The auditory cues (i.e., the spoken names of the snacks) were played upon the onset of the feedback. Half of the auditory cues were then re-played during the following NREM sleep to reactivate the social learning memories (i.e., the snack-peers’ evaluation associations). This resulted in six experimental conditions (Higher_Cued vs. Uncued; Lower_Cued vs. Uncued; Consistent_Cued vs. Uncued).

During the first lab visit, participants arrived at the lab at around 20:00. After cleaning up and the EEG setup, participants completed the Interpersonal Reactivity Index (IRI, Davis, 1983), the Socially Desirable Responding (SDR, Paulhus, 1984), and provided demographical information. Participants completed the following tasks in order. First, participants completed a psychomotor vigilance task (PVT, to measure alertness), a cue familiarization task (to get familiar with auditory cues and snack images), and an evaluation task (to indicate their baseline preferences for snacks). Second, participants performed a social learning task in which they learned about their peers’ evaluation of snacks (i.e., snack-peers’ rating associations) while hearing the spoken names of the snacks (i.e., memory reminders). Following the social learning task, participants completed the following post-learning tests: an affect misattribution procedure (AMP) task (to measure spontaneous evaluation), a speeded choice task (to measure choice), another evaluation task, and a cued recall task (to measure memories for peers’ ratings). Upon finishing these tasks, participants went to the overnight sleep session, wherein trained experimenters administered the TMR during NREM sleep.

After approximately eight hours of bedtime (12 a.m. to 8 a.m.), participants woke up and had breakfast. After ∼20 minutes of refreshing up, participants’ vigilance levels were assessed again, followed by AMP, speeded choice task, evaluation task, and cued recall task. Three days later, participants returned to the same lab and completed the same set of tasks.

#### Psychomotor vigilance task (PVT)

To test whether vigilance levels might differ across phases, participants completed a 5-minute PVT at the beginning of each phase. During the PVT, a fixation was first presented on the center of the screen with a jitter duration of 2-10 seconds. Next, a counter starting from 0 would replace the fixation. Participants shall press the button as soon as they detect the changes. Their response times (RTs) were presented on the screen as the performance feedback. We found no significant RT differences across phases, *F* (1.62, 53.41) = 1.78, *p* = .183, 𝜂^2^ = .012, suggesting no significant differences in vigilance levels across phases.

#### Cue Familiarization Task

Following the PVT, participants were familiarized with the spoken names of the snacks in the cue familiarization task. Each trial started with a 0.3 s fixation, followed by a snack image (see Figure 1 for examples), which was presented on the center of the screen for 2 s, accompanied by its spoken name (i.e., “Combos”) being played via an external speaker. The inter-trial interval (ITI) was 1 s. The task included three blocks, each containing all 48 snacks being randomly presented.

#### Evaluation Task

To assess participants’ evaluation of the snacks, we asked participants to rate their preference and familiarity with all 48 snacks four times: at pre-learning baseline, post-learning, post-TMR, and 3-day delayed phases (Figure 1B). In the evaluation task, each trial began with a 0.3 s fixation, followed by the presentation of a snack image on the screen. Using a blue triangle presented on the screen, participants then evaluated their preference for the item on a 1-11 scale (1 = Extremely Unwanted, 11 = Extremely Wanted) and their familiarity with the item (1 = Extremely Unfamiliar, 11 = Extremely Familiar). Next, we calculated the evaluation changes (ΔEvaluation) as our outcome measures by subtracting the rating between every two phases: social learning effect: post-learning minus pre-learning; TMR effect: post-TMR minus post-learning; Delayed effect: delayed minus post-learning (Figure 1A).

#### Social Learning Task

During the social learning task, participants learned their peers’ evaluations (Figure 1C). Participants were informed that their peers were students from the same university. The learning included 240 trials in 5 blocks, each containing all 48 snacks. Each trial started with a blank screen (1.2 ∼ 1.8 s), followed by a fixation cross (0.5 s). The snack image was then presented in the center of the screen for 1.5 s, together with participants’ baseline evaluation as indicated by a triangle on the preference rating scale. The scale disappeared on the screen, leaving the same snack image on the screen for 1.5 s as a buffer. Afterward, the peer’s rating was indicated by a circle on the same preference rating scale for 3 s, while the spoken name of the snack was aurally played (∼1 s) to be linked with the peers’ preference ratings. Following a 1.5 s blank screen, with only snack images being presented on the screen, participants rated the preference again (3 s maximum) using the mouse. Note that the peer ratings feedback was pre-programmed for each participant: feedback was either consistent, higher, or lower than participants’ pre-learning baseline ratings. In the higher or lower conditions, the group ratings would be 1, 2, or 3 points above or below the participants’ initial ratings, respectively. To increase the authenticity of the feedback, the chance of 3-point difference feedback was half of the probability of receiving 1 or 2-point difference feedback. We divided 48 snacks into the six experimental conditions to ensure the baseline preference ratings were comparable across conditions (for details, see Stimuli).

#### Affect Misattribution Procedure (AMP) Task

To measure the implicit evaluation for snacks, we performed the AMP task (Payne & Lundberg, 2014) in the post-learning, post-TMR, and delayed tests. Each trial of the AMP task started with a 0.3 s fixation, followed by a snack image serving as a prime. The snack image was shortly presented for 75 ms, followed by a 925 ms blank screen. Afterward, a Tibetan character was presented on the screen for 0.1 s and replaced by a mosaic image as a mask. Participants decided as soon as possible whether the target character was pleasant (“A”) or unpleasant (“L”). The AMP task contained six blocks. Forty-eight snacks were randomly presented in each block. We then calculated the update of implicit evaluation (ΔImplicit evaluation) by subtracting the percentage of choosing “pleasant” between post-TMR/delayed and post-learning phases at the item level.

#### Speeded Choice Task

Participants made speeded choices (purchase or not) toward the snacks using their own compensation in the speeded choice task. Participants completed this task three times: in the post-learning, post-TMR, and delayed tests. Each trial started with a 0.3 fixation, followed by a snack image presented on the screen for 1.5 s maximum. Participants were required to respond as soon as possible whether they would like to purchase the snack or not (“A” for yes, “L” for no). The speeded choice task contains three blocks, with 48 snacks randomly presented in each block. We then calculated the choice updating (Δ%Choose) by subtracting the percentage of choosing “Yes” between post-TMR/delayed and post-learning phases at the item level.

#### Cued Recall Task

To measure participants’ memory of their peers’ ratings for each snack, we asked them to recall and indicate their peers’ ratings in the post-learning, post-TMR, and delayed phases. In the post-learning tests, the cued recall task contained two blocks: a test with feedback block and a test without feedback block. In the feedback block, each trial began with a 0.3 s fixation, followed by a snack image and a preference rating scale being visually presented, accompanied by the spoken name of the snack. Participants clicked on the scale to indicate their peers’ preference rating. Following a 1 s blank screen, the correct ratings were presented as feedback, together with the same snack image accompanied by its spoken name aurally played. In the no-feedback block, trials were similar to those in the feedback block, except no feedback was presented. In both the post-TMR immediate and delayed phases, participants indicated their memories of peers’ ratings for each snack without feedback.

Memory error was defined as the absolute difference between participants’ recall of the feedback and the presented feedback rating. We also coded participants’ memory accuracy as follows: If participants’ recollection of peers’ ratings aligned with the feedback directions (e.g., higher, lower, consistent), the memory was deemed correct. Conversely, the memory was deemed incorrect. Thus, accuracy was coded regardless of the numerical discrepancies between the peers’ ratings and the recall.

### TMR during NREM sleep

Half of the spoken names of the snacks (24 out of 48, e.g., “Combos”) and eight additional spoken names of food items (e.g., “Celery”) were played during the TMR. These eight stimuli were never presented before the TMR and were not paired with any peers’ ratings, thus serving as non-memory control cues. Throughout the night, pink noise was played as the background noise. Well-trained experimenters monitored the EEG brainwaves and identified the sleeping stages for TMR administration. For online sleep monitoring, F3/F4, C3/C4, P3/P4, O1/O2, EOG, and EMG, with online reference at CPz, were selected. Upon detection of stable slow-wave sleep for at least 5 minutes, the names of the snacks were played via a loudspeaker placed above the participant’s head. In each block of the TMR, all 32 cues (24 snack cues and eight control cues) were randomly played (∼1 s) with an inter-stimulus interval (ISI) of 4 s. A 30 s interval separated each round of playing. The TMR phase was terminated when 20 cueing rounds were completed or reached 2 a.m., whichever came first. Cueing was stopped immediately when participants showed signs of micro-arousal or awakening and entered N1 or REM sleep. Cueing would be resumed when participants returned to stable slow-wave sleep. Participants were excluded if they received fewer than 4 TMR rounds (*n* = 9).

### EEG Acquisition

Continuous EEGs were recorded with an eego amplifier and a 64-channel gel-based waveguard cap based on an extended 10–20 layout (ANT Neuro, Enschede, and Netherlands). The online sampling rate was 500 Hz, with CPz as the online reference and AFz as the ground electrode. The horizontal electrooculogram (EOG) was recorded from an electrode placed 1.5 cm to the left external canthus. The impedance of all electrodes was maintained below 20 kΩ during the recording. During sleep, two additional electrodes were attached to both sides of the chins to measure electromyography (EMG) with a bipolar reference.

### EEG Preprocessing

Sleep EEG was processed offline using custom Python (3.8.8) scripts and MNE-Python (0.23.4) (Gramfort et al., 2013). To facilitate subsequent EEG preprocessing and analyses, the overnight EEG was cropped from 300 s ahead of the first and 300 s after the last TMR cue. Unused channels (EOG, M1, and M2) were removed from the cropped EEG data. Cropped raw EEG was filtered with a bandpass filter of 0.5-40 Hz and was notch-filtered at 50 Hz. Afterward, the EEG was downsampled to 250 Hz. Bad channels were then visually detected, removed, and interpolated. The EEG data were next re-referenced to the whole-brain average, followed by segmentation into [−15 s to 15 s] epochs relative to the onset of the cue. Bad epochs were then visually detected and removed from further analyses. Artifacts-free EEG data were further segmented into [−2 s to 6 s] epochs for time-frequency analysis. The number of remaining epochs for each condition is provided in Table S2. The overnight continuous EEG data were also retained for sleep staging and overnight spindle detection.

### Time-frequency analysis

For the time-frequency analysis, we focused on nine fronto-central channels (F1/2, Fz, FC1/2, FCz, C1/2, Cz) in accordance with recent studies examining auditory processing during sleep (Xia, Yao, et al., 2023; Züst et al., 2019)). Morlet wavelets transformation with variance cycles (three cycles at 1 Hz in length, increasing linearly with frequency to 15 cycles at 30 Hz) was applied to the [−2 s to 6 s] epochs to compute time-frequency representation (TFR) for the 1-30 Hz EEG. Next, epochs were further segmented into [−1s to 4s] epochs to eliminate edge artifacts. The trial-level spectral power was normalized (Z-scored) using [−1 s to −0.2 s] baseline of the averaged spectral power of all trials.

### Offline Automated Sleep Staging

The offline sleep staging was conducted with the YASA toolbox (0.6.1) (Vallat & Walker, 2021) implemented in Python (3.8.8). Raw overnight continuous EEG data were re-referenced to FPz according to the YASA recommendation. Sleep staging was based on C4 (or C3 if C4 was marked as a bad channel) and EOG (see Table S3 for sleep stage information).

### Spindle Detection

The automated spindle detection was implemented in the YASA toolbox (0.6.1) (Vallat & Walker, 2021). The spindle detection algorithm was applied separately to the preprocessed overnight continuous EEG data and artifacts-free [−15 s to 15 s] epochs. We applied three thresholds in identifying a spindle: 1) relative power, which indicated the power in the sigma frequency range (11-16 Hz) relative to the total power in the broadband frequency (1-30 Hz), 2) correlation, the correlation between sigma-filtered signal and broadband signal, and 3) RMS, moving root mean square (RMS) of the sigma-filtered signal. Overnight spindle detection was applied on the continuous preprocessed EEG data at the Cz during N2 (relative power = 0.2, correlation = 0.65, RMS = 1.5) and N3 (relative power = None, correlation = 0.50, RMS = 1.5) sleep stages separately. We adopted different parameters for the N2 and N3 sleep stages because they showed distinct EEG characteristics. Spindle density was then calculated using the following formula:

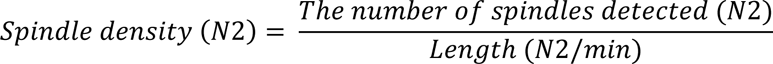

The spindle detection algorithm was also applied to the artifacts-free segmented data, with the same parameters as the overnight spindle detection of the N3 sleep stage, as most of our TMR cues were played during the N3 sleep. Subsequently, the algorithm generated a series of binary values (spindle presence or absence) to indicate whether a spindle was detected at each timepoint (each timepoint represented 4 ms). The cue-elicited spindle probability was next determined by computing the proportion of detected spindle across trials at each timepoint (Schechtman et al., 2021; Xia, Yao, et al., 2023).

### Statistical Analysis

First, we investigated the impact of social learning and TMR on changes in evaluation, implicit evaluation, speeded choice, and memory error. We conducted repeated-measure ANOVA with R (4.2.2) and the afex package (1.2.1) implemented in R. We further examined the effects of social learning, TMR, and subsequent memory on evaluation changes. Due to the limited number of trials after separating trials into correctly vs. incorrectly remembered, we adopted an item-level linear mixed model. To deal with the singular fitting problem, we chose a Bayesian linear mixed model (BLMM) with R using the brms package (2.20.4) (Bürkner, 2021). Since evaluations were only tested once in each phase, the evaluation changes at the item level are discrete (from −8 to 8). Therefore, we adopted a cumulative distribution in the BLMM and transformed the evaluation changes into ordinal-level data. The following BLMM was applied:

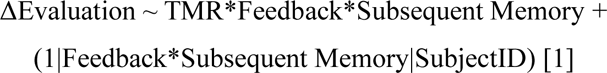

Next, we investigated whether cues would elicit significantly different EEG power changes and spindle probability. We employed a cluster-based two-tailed one-sample permutation test, implemented in the MNE toolbox with 1000 randomizations and a statistical threshold of 0.05.

To quantify the relationship between cue-elicited power and evaluation changes, we continued to utilize item-level BLMM. The cue-elicited power was extracted from the significant clusters at the item level. We also adopted a cumulative distribution and transformed the evaluation changes to ordinal-level data. Because we considered that the cueing repetition could impact the signal- to-noise ratio of EEG data, we took the repetition number (N) as a control variable. The following BLMM was employed:

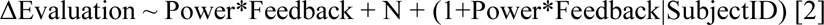

The same item-level BLMM was employed to investigate the relationship between cue-elicited spindle probability and evaluation changes:

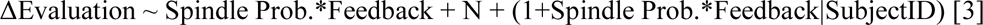

We were also interested in the impact of overnight spindle density on the evaluation changes. The following subject-level BLMMs were utilized for cued and uncued snacks, respectively:

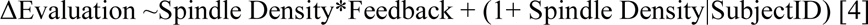

Statistical inferences for the BLMM were based on the 95% highest density interval (HDI) of the posterior distribution. Effects were considered significant if the 95% HDI did not encompass 0. Note that we focused on performance from higher and lower conditions, wherein participants were expected to change their evaluations.

## Results

### Effects of social learning and TMR on evaluation changes

We began by examining whether social learning modulated evaluations of the snacks. In a TMR (cued vs. uncued) by feedback (higher vs. lower) repeated measure ANOVA, we found the expected social learning effect: feedback significantly modulated ΔEvaluation (i.e., changes of evaluation from pre- to post-learning; *F* (1, 33) = 23.42, *p* < .001, 𝜂^2^ = 0.18; Figure 2A). Specifically, when peers’ evaluations were higher than participants’ initial evaluations, participants’ evaluations increased accordingly. In contrast, the TMR effect was not significant (*F* (1, 33) = 0.02, *p* = .877, 𝜂^2^ < 0.01) nor was the TMR by feedback interaction (*F* (1, 33) = 0.34, *p* = .564, 𝜂^2^ < 0.01), indicating that cued and uncued snacks showed comparable social learning effects before sleep and TMR manipulation.

**Figure 2:**
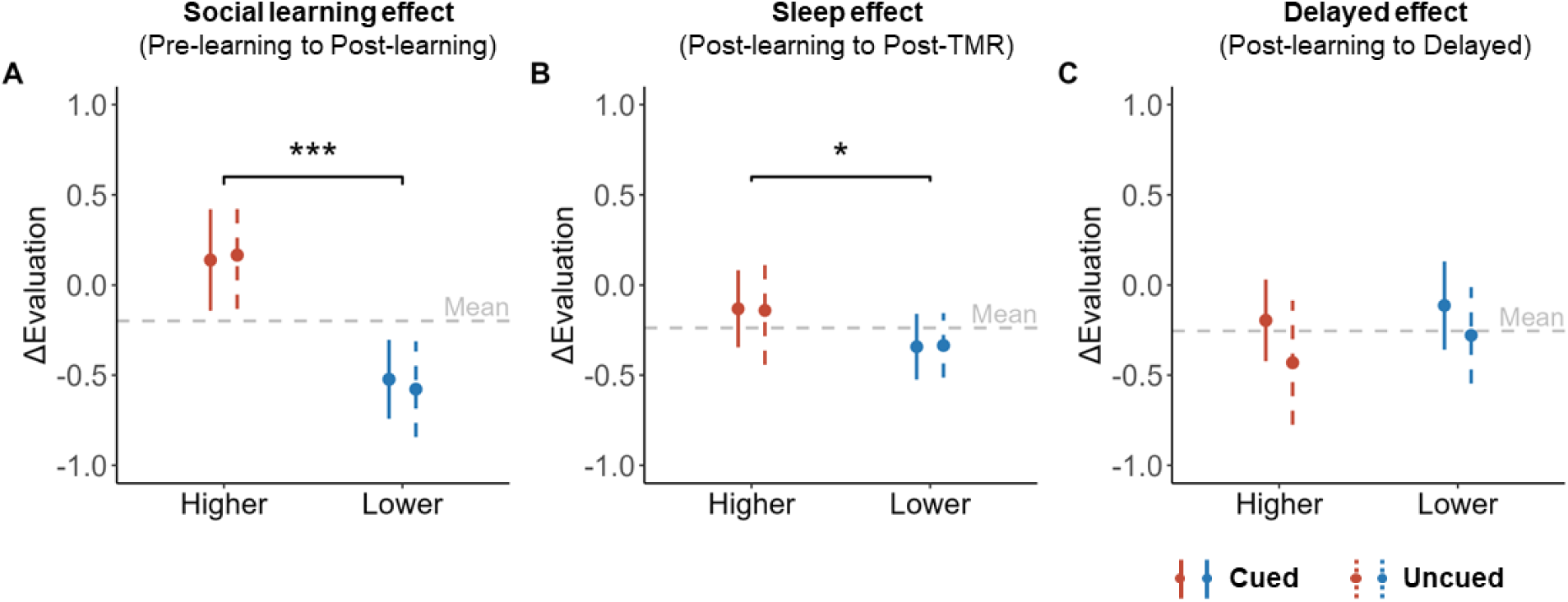
Effects of feedback (i.e., peers’ ratings either higher or lower than pre-learning baseline ratings) and TMR (cued vs. uncued) on ΔEvaluation from (A) pre-learning to post-learning, (B) post-learning to post-TMR, and (C) post-learning to delayed phases. The error bars indicated the 95% confidence intervals. The horizontal grey dashed line represents the mean of ΔEvaluation at the corresponding phase. ***: *p* < .001. *: *p* < .05.

We next examined the impact of sleep TMR on the ΔEvaluation from the post-learning to post-TMR phase. We again found a significant main effect of feedback, such that the ΔEvaluation was significantly increased for the higher than for the lower feedback condition (*F* (1, 33) = 4.72, *p* = .037, 𝜂^2^ = 0.03; Figure 2B). This significant feedback effect on ΔEvaluation indicated that the difference between higher vs. lower feedback directions further enlarged from post-learning to post-TMR phases. Contrary to our hypotheses, neither the TMR (cued vs. uncued) nor the TMR by feedback interaction was significant (*F* (1, 33) < 0.01, *p* = .994, 𝜂^2^ < 0.01; *F* (1, 33) = 0.01, *p* = .911, 𝜂^2^ < 0.01, respectively).

We further examined the 3-day delay effect of sleep TMR on the ΔEvaluation from post-learning to the 3-day delayed phase. We found a non-significant trend of the TMR effect: cued snacks showed numerically higher ΔEvaluation than uncued snacks (*F* (1, 33) = 3.69, *p* = .063, 𝜂^2^ = 0.02; Figure 2C). However, neither feedback (*F* (1, 33) = 1.23, *p* = .275, 𝜂^2^ = 0.01) nor interaction effects (*F* (1, 33) = 0.18, *p* = .677, 𝜂^2^ < 0.01) were significant. We postulated that the cueing might increase familiarity, thus enhancing preferences (see Ai et al., 2018). Indeed, in a TMR by feedback repeated measure ANOVA on the familiarity rating, we found that cueing significantly enhanced familiarity ratings of snacks in the 3-day delayed session (*F* (1, 33) = 8.28, *p* = .007, 𝜂^2^ = 0.03), but not in the post-learning nor post-TMR tests (*p*s > .116). Thus, the numerically higher evaluations of cued snacks could be attributed to their higher familiarity at the delayed phase.

### Effects of social learning and TMR on memory errors

Here, we examined whether TMR changed memory errors, i.e., the absolute numerical differences between participants’ recalled peers’ ratings and the presented peers’ ratings. In the TMR by feedback repeated measure ANOVA, we did not find a significant main or interaction effect in the post-learning phase (*p*s > .487). In the post-TMR phase, we observed a non-significant trend of increased memory error for the higher than the lower feedback conditions (*F* (1, 33) = 4.01, *p* = .054, 𝜂^2^ = 0.02). However, no significant main effect of TMR (*F* (1, 33) = 0.96, *p* = .333, 𝜂^2^_𝐺_ < 0.01), and the interaction effect was observed (*F* (1, 33) = 0.02, *p* = .879, 𝜂^2^_𝐺_ < 0.01). In the delayed phase, no significant main effects nor interaction effects were found (*p*s > .230).

### Relationship between subsequent memory accuracies and evaluation changes

Although TMR did not influence memory errors when recalling peers’ ratings, we examined whether evaluation changes were associated with memory accuracies, i.e., whether participants’ recall of the peers’ ratings aligned with the feedback directions. Therefore, we conducted feedback by TMR by subsequent memory (correctly vs. incorrectly remembered) three-way item-level BLMM for ΔEvaluation.

For the ΔEvaluation from pre-learning to post-learning, we found a significant interaction between subsequent memory and feedback (median = 2.93, 95% HDI [1.93, 3.85], Figure 3A). Post-hoc analysis revealed that when participants correctly remembered the feedback direction, the ΔEvaluation in the higher feedback condition was significantly higher than that in the lower feedback condition (higher vs. lower, median*_diff_* = 1.55, 95% HDI [1.14, 1.98]). Conversely, when participants incorrectly remembered the feedback direction, the ΔEvaluation in the higher feedback condition was significantly lower than in the lower feedback condition (median*_diff_* = −1.54, 95% HDI [−2.18, −0.85]). The other main and interaction effects were insignificant (−1.52< median <0.32).

**Figure 3:**
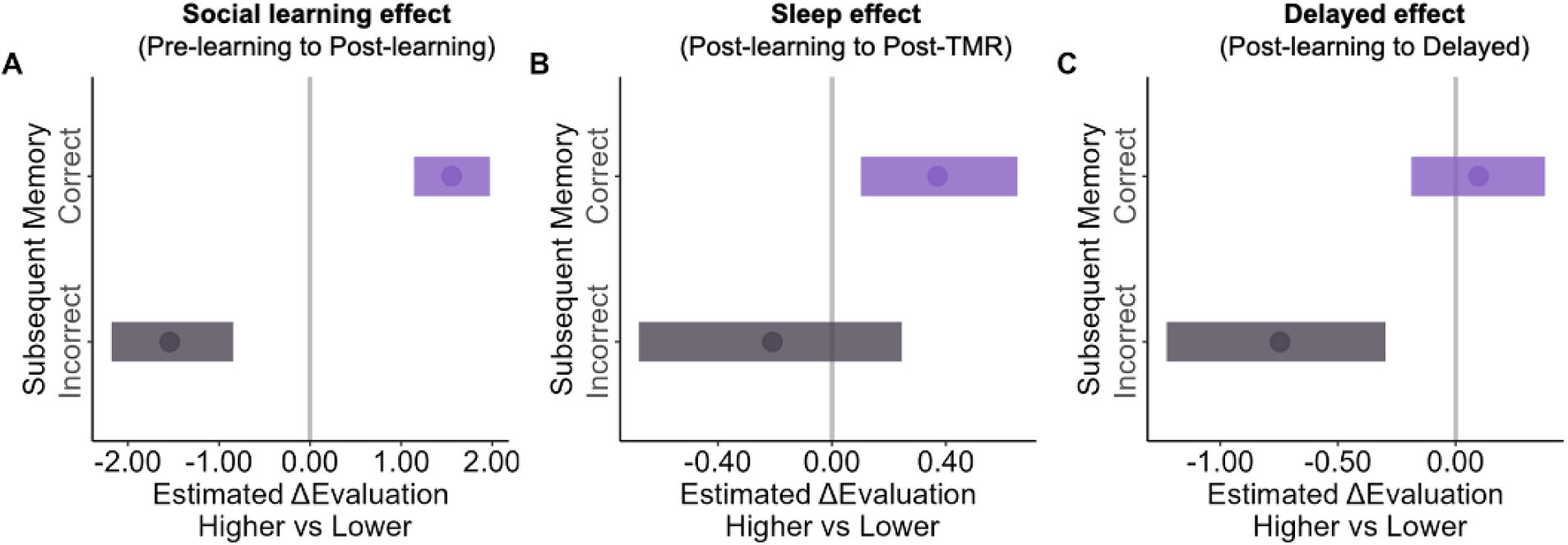
Effects of subsequent memory, TMR, and feedback on ΔEvaluation from (A) pre-learning to post-learning, (B) post-learning to post-TMR, and (C) post-learning to delayed phases. The horizontal lines indicated the 95% highest density interval (HDI), and vertical gray lines correspond to 0. The dot indicated the median point. If the 95% HDI did not encompass 0, the result would be considered significant.

For the ΔEvaluation from post-learning to post-TMR, we similarly found a significant subsequent memory by feedback interaction effect (median = 0.74, 95% HDI [0.03, 1.46], Figure 3B). Post-hoc analyses revealed that when participants correctly remembered the feedback direction, the ΔEvaluation in the higher feedback condition was significantly higher than that in the lower condition (median*_diff_* = 0.37, 95% HDI [0.10, 0.65]). In contrast, when participants incorrectly remembered the feedback direction, the ΔEvaluation did not differ between the higher and the lower condition (median*_diff_* = −0.21, 95% HDI [−0.68, 0.25]).

For the ΔEvaluation from post-learning to the delayed phase, the same BLMM again revealed a significant interaction effect (median = 0.71, 95% HDI [0.01, 1.40], Figure 3C). Post-hoc analyses revealed that when participants correctly remembered the feedback direction, the ΔEvaluation between the higher and the lower condition did not significantly differ (median*_diff_* = 0.10, 95% HDI [−0.19, 0.38]). In contrast, when participants incorrectly remembered the feedback direction, the ΔEvaluation of the higher condition was significantly lower than that in the lower condition (median*_diff_* = −0.75, 95% HDI [−1.23, −0.30]). These results suggested that the evaluation changes were related to the memory of the feedback directions across all three phases.

### Effects of social learning and TMR on implicit evaluation and speeded choice

Observing the social learning effects on subjective evaluation changes, we further examined whether social learning and TMR could impact implicit evaluation (ΔImplicit evaluation based on AMP performance) and speeded choices (Δ%Choose based on the speeded choice task) by conducting TMR by feedback repeated measure ANOVAs.

In the speeded choice task, we observed a significant main effect of feedback in Δ%Choose from post-learning to post-TMR phases, that participants were more willing to choose the snacks in the higher than the lower feedback conditions (*F* (1, 32) = 4.83, *p* = .035, 𝜂^2^ = 0.03). No significant effect of TMR nor their interaction was found (*p*s >.316; Figure S1A). Similarly, no significant effect of feedback, TMR, nor their interaction in Δ%Choose from post-learning to delayed phases was observed (*p*s > .283; Figure S1B).

In the AMP, we did not observe a significant effect of feedback, TMR, nor their interaction in the ΔImplicit evaluation from post-learning to post-TMR (*p*s > .312) and to delayed phases (*p*s > .398) (Figure S1C-D).

### Cue-elicited delta-theta power predicted evaluation changes of cued snacks

Even though we did not observe the TMR effect on Δevaluation during the post-TMR phases, we proceeded to perform sleep EEG analyses to investigate the neural mechanism that could drive the overall enhanced social learning effect for both cued and uncued snacks.

We first examined whether presenting cues during sleep would elicit significant EEG power changes relative to the pre-cue baseline (i.e., −1000 to −200 ms prior to the cue onset). We found that the cues significantly enhanced the 1-30 Hz power during an early cluster (−96 to 2928 ms relative to the cue onset, *p_cluster_* = .001, corrected for multiple comparisons by cluster-based permutation test; see Methods) but reduced the 5.5 – 18.5 Hz power in a later cluster (2132 to 4000 ms, *p_cluster_* = .025, Figure 4A). However, we did not find significant EEG power differences between the higher and lower feedback conditions (*p_cluster_*s > .085, Figure S2B-D). Similarly, the control cues enhanced the 1-30 Hz EEG power in the early cluster (−-360 to 3028 ms relative to the cue onset, *p_cluster_* = .001) but reduced the 8.5 to 17.5 Hz power in the later cluster (2136 to 4000 ms, *p_cluster_* = .047, Figure 4B). However, further analysis did not reveal significant EEG differences between memory and control cues (*p_cluster_*s > .217, Figure S2A). These EEG power changes suggested that both memory and control cues were processed during sleep.

**Figure 4:**
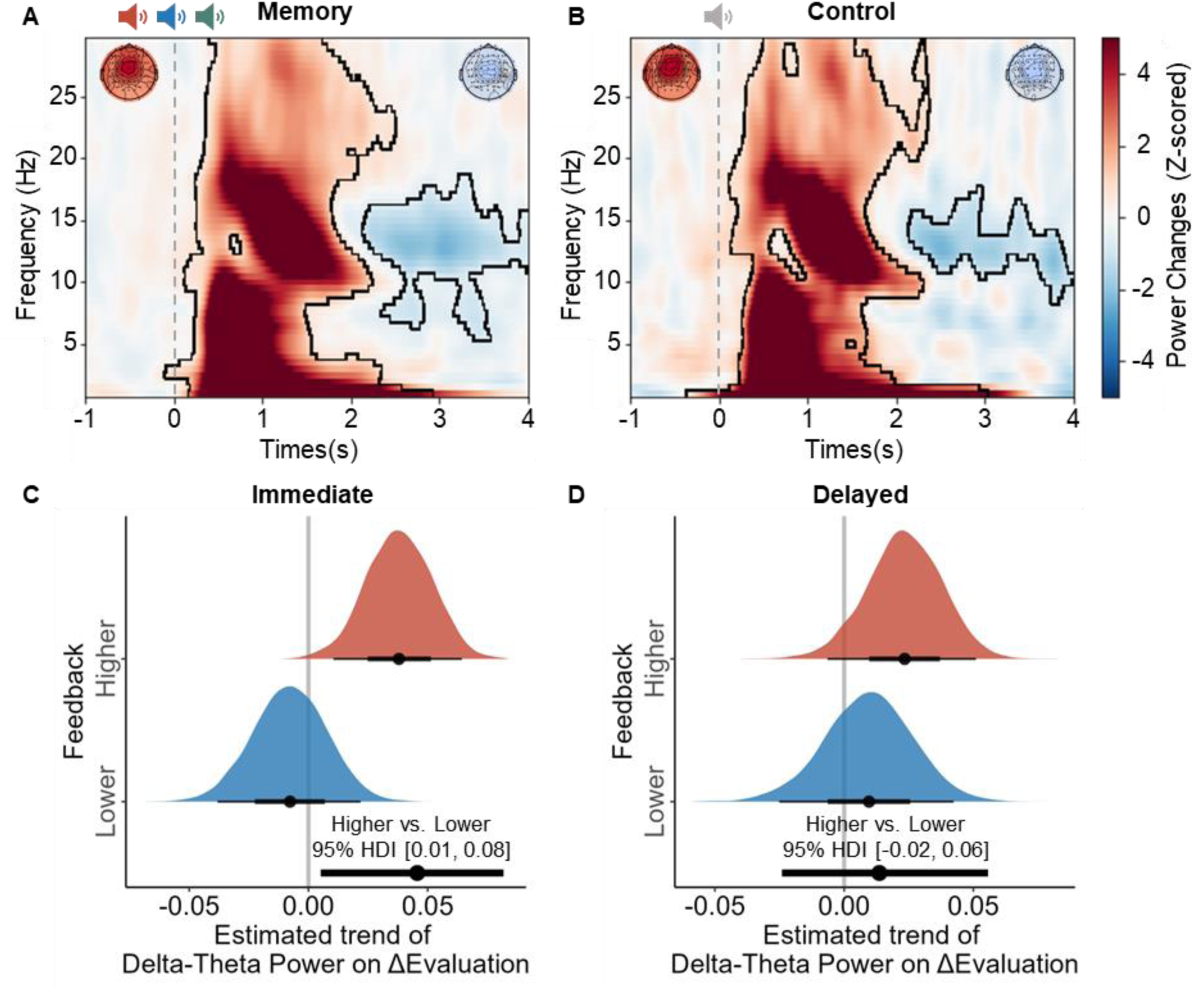
Cue-elicited EEG Power and ΔEvaluataion. A) Memory cue (higher, lower, and consistent) and B) control cue-elicited power spectral. The topography on the left-top and right-top corners indicated the power at each channel at the early and late clusters, respectively. The contour highlighted significant clusters. The effect of memory cue-elicited delta-theta power (1-8 Hz) on ΔEvaluation of cued snacks from C) post-learning to post-TMR and D) post-learning to delayed phases. The black line below the red and blue density plot indicated the 95% highest density interval (HDI) for higher and lower feedback conditions respectively. The bottom black line indicates the difference higher vs. lower feedback conditions. The dot indicated the median point. If the 95% HDI did not encompass 0, the result would be considered significant.

We next examined whether memory cue-elicited EEG power could predict the Δevaluation of cued snacks by employing the BLMM. We extracted cue-elicited delta-theta power (1-8 Hz) and sigma power (12-16 Hz) within the early identified cluster and the 0-2 s at the item level. We selected the 0-2 s because this time window captured the early cluster yet did not overlap with the late cluster. The EEG power by feedback BLMM showed a significant interaction (higher vs. lower, median*_diff_* = 0.05, 95% HDI [0.01, 0.08], Figure 4C), such that the cue-elicited delta-theta power predicted the post-TMR immediate evaluation changes for cued snacks as a function of feedback (Δevaluation from post-learning to post-TMR phase). Post-hoc analyses showed a significant positive prediction of delta-theta power for Δevaluation (median = 0.04, 95% HDI [0.01, 0.06]) in the higher feedback condition, but not in the lower feedback condition (median = −0.01, 95% HDI [−0.04, 0.02]). This result indicated that the higher the cue-elicited delta-theta power, the larger the changes in evaluations were in the higher feedback condition compared to the lower feedback condition.

We further examined whether cue-elicited delta-theta power predicted delayed ΔEvaluation. However, no significant interaction effects were found (higher vs. lower, median*_diff_* = 0.01, 95% HDI [−0.02, 0.06], Figure 4D). Additionally, we did not observe significant effects of cue-elicited sigma power in either immediate (median*_diff_* = 0.04, 95% HDI [−0.02, 0.09]) and delayed ΔEvaluation (median*_diff_* = −0.00, 95% HDI [−0.06, 0.06]).

### Overnight N2 sleep spindle density predicted evaluation changes for cued snacks

Given the sleep spindle’s crucial role in sleep-mediated memory consolidation (Antony et al., 2019), we further examined the relationship between cued-elicited and overnight spindle activities and the evaluation changes.

First, for cue-elicited spindle activity, we found that compared to the pre-stimulus baseline, both memory (*p*_cluster_ = .003) and control cues (*p*_cluster_ = .016) elicited significantly higher spindle probabilities (Figure S3A) approximately 1 second after the cue onset. However, there was no significant difference between memory vs. control cue-elicited spindle probability, nor among the different feedback conditions within memory cues (cluster-based permutation tests, *p*_cluster_s > .497, Figure S3B). Furthermore, the cue-elicited spindle probability did not predict ΔEvaluation for the cued snack in both the immediate (median*_diff_* = 2.10, 95% HDI [−3.49, 7.50]) and the delayed phase (median*_diff_* = −2.03, 95% HDI [−7.81, 3.45]).

Next, we investigated the relationship between overnight spindle density and ΔEvaluation for cued and uncued snacks separately. For the cued snacks, the subject-level BLMM revealed that the overnight N2 spindle density predicted overnight ΔEvaluation (from post-learning to post-TMR), as indicated by the significant spindle density by feedback interaction (higher vs. lower, median*_diff_* = 0.17, 95% HDI [0.03, 0.31], Figure 5A). That is, higher overnight spindle density was associated with increased evaluation changes for the higher feedback condition than the lower feedback condition.

**Figure 5:**
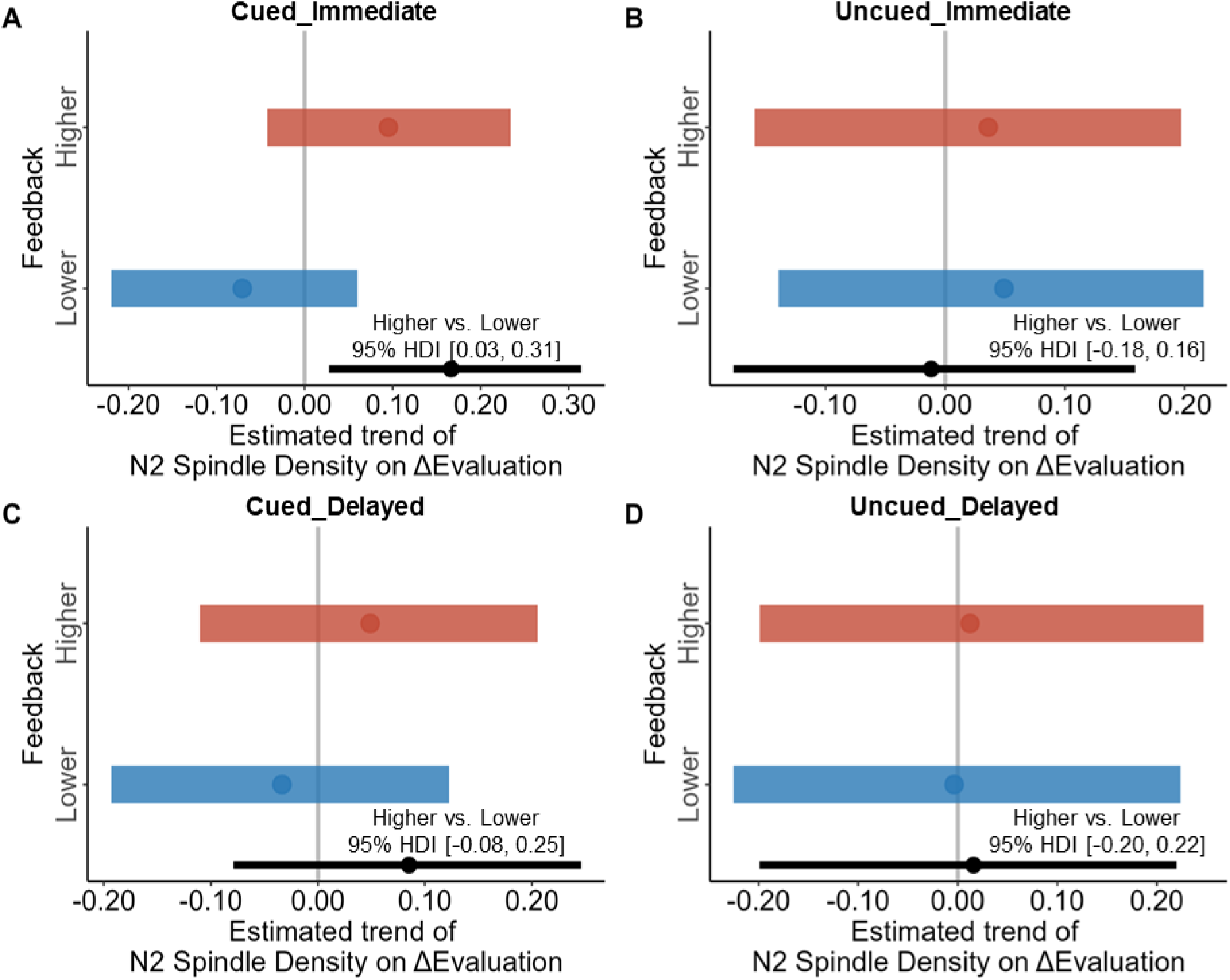
The relationship between overnight N2 Spindle Density and ΔEvaluation of cued snacks or uncued snacks from post-learning to post-TMR phases (A, B) and from post-learning to delayed phases (C, D). The vertical gray lines correspond to 0. The horizontal red and blue lines indicated the 95% highest density interval (HDI) for higher and lower feedback conditions respectively. The bottom black line indicates the difference higher vs. lower feedback conditions. The dot indicated the median point. If the 95% HDI did not encompass 0, the result would be considered as significant.

Again, no such effects were observed in the 3-day delayed test (median*_diff_* = 0.09, 95% HDI [−0.08, 0.25], Figure 5C), nor were observed for uncued snacks in either overnight or delayed ΔEvaluation (median*_diff_* = −0.01, 95% HDI [−0.18, 0.16], Figure 5B; median*_diff_* = 0.02, 95% HDI [−0.20, 0.22], Figure 5D).

## Discussion

People often change their evaluations and opinions upon learning about their peers’ evaluations and choices, i.e., social learning (Berns et al., 2010; Campbell-Meiklejohn et al., 2010; Kendal et al., 2018). Moreover, sleep impacts social and non-social decision-making (Ben Simon et al., 2022; Dickinson & McElroy, 2017; Holbein et al., 2019; Venkatraman et al., 2011). Combining the social learning paradigm with sleep-based targeted memory reactivation (TMR), we investigated whether reactivating the daytime social learning experience during non-rapid-eye-movement (NREM) sleep could further promote social learning-induced evaluation changes. We found that the social learning-induced evaluation changes enlarged following one night of sleep, though TMR did not selectively enhance these changes. Despite the lack of cueing effect, examining sleep EEG activity showed that the cue-elicited delta-theta (1-8 Hz) EEG power and the overnight N2 spindle density predicted the overnight evaluation changes of cued snacks. Together, we provided new evidence that the sleep-mediated memory reactivation processes could fortify evaluation changes induced by social learning.

TMR has been shown to benefit various types of learning by promoting sleep-mediated memory consolidation (Hu et al., 2020). However, research on TMR’s impact on social learning is limited. A previous study endeavoured to examine how TMR influences interpersonal trust yet reported no significant sleep nor TMR effect (Strachan et al., 2020). Although we did not find a significant TMR effect in the post-TMR immediate test, it was noteworthy that sleep EEG activity related to memory reactivation facilitated overnight evaluation changes for cued but not uncued snacks. Specifically, for cued snacks, we found that both cue-elicited delta-theta power and the overnight N2 spindle density differentially predicted evaluation changes between the higher and lower feedback conditions. Mounting evidence suggests that these two EEG features characterize cue-elicited and spontaneous memory reactivation during sleep, respectively (Clemens et al., 2005; Mednick et al., 2013; Petzka et al., 2022; Schönauer et al., 2017; Schreiner et al., 2021). Specifically, these findings are consistent with previous TMR studies that also demonstrated the beneficial role of cue-elicited delta-theta power in evaluation updates (Ai et al., 2018; Xia, Antony, et al., 2023) and long-term memory maintenance (Liu et al., 2023; Oudiette et al., 2013; Rihm et al., 2014). Moreover, sleep spindles are instrumental to memory re-processing during sleep (Antony et al., 2019; Petzka et al., 2022), such that category- and even item-specific neural representation could be evident during cue-elicited spindle activity (Cairney et al., 2018; Liu et al., 2023). A previous TMR study also found that the spindle density during a nap supported the TMR cueing benefits (Creery et al., 2015). Our results further suggest that the sleep spindles could support overnight evaluation changes implicating social learning, presumably via memory reactivation-related processes.

Our findings contribute to the theoretical understanding of how memory-related processing impacts evaluations in social learning and sleep (Amodio, 2019; Biderman et al., 2020). We found that only when participants could correctly remember the feedback direction they showed the social learning effect by following peers’ evaluations. Contrary to previous research that focused on memory interference that weakens memories (Biderman et al., 2023), our study aimed to change evaluation by promoting memories through sleep-mediated memory reactivation. Building on previous TMR and sleep research that aims to enhance evaluative memory or familiarity to update evaluations (Ai et al., 2018; Jin et al., 2023), we further showed that TMR and overnight sleep influenced social learning-induced evaluation changes.

In addition to memory accuracies that capture episodic retrieval of peers’ evaluations, we also measured participants’ familiarity ratings towards the snacks. Intriguingly, we found that TMR increased familiarity with the cued snacks in the 3-day delayed session, which may influence the delayed evaluations. This finding aligned with well-established findings that people preferred familiar over unfamiliar snacks (Aldridge et al., 2009; Raudenbush & Frank, 1999) and the findings that merely re-playing snacks’ names during sleep could enhance people’s preference toward these snacks (Ai et al., 2018). Notably, the TMR’s benefits in strengthening familiarity emerged in the delayed but not in the immediate test. This finding is consistent with recent research showing that TMR often showed delayed benefits in memory performance (Cairney et al., 2018; Rakowska et al., 2021). One intriguing question that warrants future research is the respective impacts of episodic memory and familiarity on human evaluations and decision-making, and how sleep may influence different retrieval processes that support decision-making.

Limitations and future directions shall be discussed. First, we did not find a significant TMR effect on evaluation changes, even though item-level cue-elicited EEG activity predicted evaluation changes for cued snacks. One possibility is that the reactivation may generalize to uncued snacks, given that cued and uncued snacks share the same learning context (Oudiette et al., 2013; Schechtman et al., 2023). Future research shall test whether and when generalization occurred. Second, the classic social learning paradigm adopted here involved passive observation of peers’ evaluations in laboratory settings. Secondly, given that social learning often happens during real-life interpersonal interactions (Pan et al., 2022; Zhang & Gläscher, 2020), future research shall examine the role of sleep and TMR in consolidating more realistic social learning experiences. Lastly, while people are intrinsically motivated to follow peers’ opinions (Klucharev et al., 2009), given the universal need to seek social belongingness (Baumeister & Leary, 1995; Izuma, 2013), our study did not manipulate extrinsic motivations involved in many social learning scenarios. For example, successful social learning can lead to social rewards, while unsuccessful learning may incur punishments (Molho et al., 2020). Given that motivational processes could bias memory reactivation during sleep (Sterpenich et al., 2021; Wilhelm et al., 2011), future research shall consider manipulating motivational processes during social learning and how sleep and memory reactivation interact with motivation to change behavior.

In conclusion, we found that the social learning-induced evaluation changes became more pronounced after sleep, irrespective of whether or not the corresponding memories were exogenously reactivated during sleep. Sleep EEG activity, such as the cue-elicited delta-theta power and the overnight N2 spindle activity, supported the evaluation changes for the cued snacks. Our research contributes to the theoretical understanding of memory-based evaluation by highlighting the significance of offline sleep-mediated memory reactivation processes. Considering social learning can influence moral decision-making (Yu et al., 2021) and healthy behavior (Bavel et al., 2020; Chung et al., 2020; Nook & Zaki, 2015; Templeton et al., 2016), using TMR and sleep in conjunction with social learning may offer insights into fostering adaptive behaviors in a social and healthy context.

## Acknowledgment

The authors thank Hinako Kojima and Tiffany Wing Lam Yip for their assistance in data collection. This research was supported by the Ministry of Science and Technology of China STI2030-Major Projects (No. 2022ZD0214100), National Natural Science Foundation of China (No. 32171056), General Research Fund (No. 17614922) of Hong Kong Research Grants Council to X. H. The funders were not involved in the study design, data collection and analysis, publication decisions, or manuscript preparation.

## Author contributions

**D.C.:** Conceptualization, Investigation, Formal Analysis, Data Curation, Software, Methodology, Writing – Original Draft, Writing – Review & Editing, Visualization; **T.X.:** Methodology, Formal Analysis, Validation, Writing – Review & Editing; **Z.Y.:** Methodology, Formal Analysis, Validation, Writing – Review & Editing; **L.Z.**: Investigation, Writing – Review & Editing; **X.H.**: Conceptualization, Writing - Original Draft, Writing - Review & Editing, Supervision, Project Administration, Funding Acquisition.

## Data availability

Preprocessed data is available on the Open Science Framework (OSF) at https://osf.io/t96z5/?view_only=8bc5488d7d854455b19fe792e693410d.

## Code availability

Statistical analysis scripts are available on the Open Science Framework (OSF) at https://osf.io/t96z5/?view_only=8bc5488d7d854455b19fe792e693410d.

## Reference

Ai, S., Yin, Y., Chen, Y., Wang, C., Sun, Y., Tang, X., Lu, L., Zhu, L., & Shi, J. (2018). Promoting subjective preferences in simple economic choices during nap. eLife, 7, e40583. 10.7554/eLife.40583

Aldridge, V., Dovey, T. M., & Halford, J. C. G. (2009). The role of familiarity in dietary development. Developmental Review, 29(1), 32–44. 10.1016/j.dr.2008.11.001

Amodio, D. M. (2019). Social Cognition 2.0: An Interactive Memory Systems Account. Trends in Cognitive Sciences, 23(1), 21–33. 10.1016/j.tics.2018.10.002

Antony, J. W., Gobel, E. W., O’Hare, J. K., Reber, P. J., & Paller, K. A. (2012). Cued memory reactivation during sleep influences skill learning. Nature Neuroscience, 15(8), Article 8. 10.1038/nn.3152

Antony, J. W., Schönauer, M., Staresina, B. P., & Cairney, S. A. (2019). Sleep Spindles and Memory Reprocessing. Trends in Neurosciences, 42(1), 1–3. 10.1016/j.tins.2018.09.012

Baumeister, R. F., & Leary, M. R. (1995). The need to belong: Desire for interpersonal attachments as a fundamental human motivation. Psychological Bulletin, 117(3), 497–529. 10.1037/0033-2909.117.3.497

Bavel, J. J. V., Baicker, K., Boggio, P. S., Capraro, V., Cichocka, A., Cikara, M., Crockett, M. J., Crum, A. J., Douglas, K. M., Druckman, J. N., Drury, J., Dube, O., Ellemers, N., Finkel, E. J., Fowler, J. H., Gelfand, M., Han, S., Haslam, S. A., Jetten, J., … Willer, R. (2020). Using social and behavioural science to support COVID-19 pandemic response. Nature Human Behaviour, 4(5), Article 5. 10.1038/s41562-020-0884-z

Ben Simon, E., Vallat, R., Rossi, A., & Walker, M. P. (2022). Sleep loss leads to the withdrawal of human helping across individuals, groups, and large-scale societies. PLOS Biology, 20(8), e3001733. 10.1371/journal.pbio.3001733

Berns, G. S., Capra, C. M., Moore, S., & Noussair, C. (2010). Neural mechanisms of the influence of popularity on adolescent ratings of music. NeuroImage, 49(3), 2687–2696. 10.1016/j.neuroimage.2009.10.070

Biderman, N., Bakkour, A., & Shohamy, D. (2020). What Are Memories For? The Hippocampus Bridges Past Experience with Future Decisions. Trends in Cognitive Sciences, 24(7), 542–556. 10.1016/j.tics.2020.04.004

Biderman, N., Gershman, S. J., & Shohamy, D. (2023). The role of memory in counterfactual valuation. Journal of Experimental Psychology: General, 152(6), 1754–1767. 10.1037/xge0001364

Brady, W. J., McLoughlin, K., Doan, T. N., & Crockett, M. J. (2021). How social learning amplifies moral outrage expression in online social networks. Science Advances, 7(33), eabe5641. 10.1126/sciadv.abe5641

Brodt, S., Inostroza, M., Niethard, N., & Born, J. (2023). Sleep—A brain-state serving systems memory consolidation. Neuron, 111(7), 1050–1075. 10.1016/j.neuron.2023.03.005

Bürkner, P.-C. (2021). Bayesian Item Response Modeling in R with brms and Stan. Journal of Statistical Software, 100, 1–54. 10.18637/jss.v100.i05

Cairney, S. A., Guttesen, A. á V., El Marj, N., & Staresina, B. P. (2018). Memory Consolidation Is Linked to Spindle-Mediated Information Processing during Sleep. Current Biology, 28(6), 948–954.e4. 10.1016/j.cub.2018.01.087

Cairney, S. A., Sobczak, J. M., Lindsay, S., & Gaskell, M. G. (2017). Mechanisms of Memory Retrieval in Slow-Wave Sleep. Sleep, 40(9). 10.1093/sleep/zsx114

Campbell-Meiklejohn, D. K., Bach, D. R., Roepstorff, A., Dolan, R. J., & Frith, C. D. (2010). How the Opinion of Others Affects Our Valuation of Objects. Current Biology, 20(13), 1165–1170. 10.1016/j.cub.2010.04.055

Chen, D., Yao, Z., Liu, J., Wu, H., & Hu, X. (2023). In-group Social Conformity Updates the Neural Representation of Facial Attractiveness. bioRxiv. 10.1101/2023.02.08.527779

Chung, D., Orloff, M. A., Lauharatanahirun, N., Chiu, P. H., & King-Casas, B. (2020). Valuation of peers’ safe choices is associated with substance-naïveté in adolescents. Proceedings of the National Academy of Sciences, 117(50), 31729–31737. 10.1073/pnas.1919111117

Clemens, Z., Fabó, D., & Halász, P. (2005). Overnight verbal memory retention correlates with the number of sleep spindles. Neuroscience, 132(2), 529–535. 10.1016/j.neuroscience.2005.01.011

Creery, J. D., Oudiette, D., Antony, J. W., & Paller, K. A. (2015). Targeted Memory Reactivation during Sleep Depends on Prior Learning. Sleep, 38(5), 755–763. 10.5665/sleep.4670

Davis, M. H. (1983). Measuring individual differences in empathy: Evidence for a multidimensional approach. Journal of Personality and Social Psychology, 44(1), 113–126. 10.1037/0022-3514.44.1.113

Dickinson, D. L., & McElroy, T. (2017). Sleep restriction and circadian effects on social decisions. European Economic Review, 97, 57–71. 10.1016/j.euroecorev.2017.05.002

Gramfort, A., Luessi, M., Larson, E., Engemann, D., Strohmeier, D., Brodbeck, C., Goj, R., Jas, M., Brooks, T., Parkkonen, L., & Hämäläinen, M. (2013). MEG and EEG data analysis with MNE-Python. Frontiers in Neuroscience, 7. https://www.frontiersin.org/articles/10.3389/fnins.2013.00267

Hare, T. A., Malmaud, J., & Rangel, A. (2011). Focusing Attention on the Health Aspects of Foods Changes Value Signals in vmPFC and Improves Dietary Choice. Journal of Neuroscience, 31(30), 11077–11087. 10.1523/JNEUROSCI.6383-10.2011

Holbein, J. B., Schafer, J. P., & Dickinson, D. L. (2019). Insufficient sleep reduces voting and ther prosocial behaviours. Nature Human Behaviour, 3(5), 492–500. 10.1038/s41562-019-0543-4

Hu, X., Antony, J. W., Creery, J. D., Vargas, I. M., Bodenhausen, G. V., & Paller, K. A. (2015). Unlearning implicit social biases during sleep. Science, 348(6238), 1013–1015. 10.1126/science.aaa3841

Hu, X., Cheng, L. Y., Chiu, M. H., & Paller, K. A. (2020). Promoting memory consolidation during sleep: A meta-analysis of targeted memory reactivation. Psychological Bulletin, 146(3), 218–244. 10.1037/bul0000223

Huang, Y., Kendrick, K. M., & Yu, R. (2014). Conformity to the Opinions of Other People Lasts for No More Than 3 Days. Psychological Science, 25(7), 1388–1393. 10.1177/0956797614532104

Humiston, G. B., & Wamsley, E. J. (2019). Unlearning implicit social biases during sleep: A failure to replicate. PLOS ONE, 14(1), e0211416. 10.1371/journal.pone.0211416

Hütter, M. (2022). An integrative review of dual- and single-process accounts of evaluative conditioning. Nature Reviews Psychology, 1(11), 640–653. 10.1038/s44159-022-00102-7

Izuma, K. (2013). The neural basis of social influence and attitude change. Current Opinion in Neurobiology, 23(3), 456–462. 10.1016/j.conb.2013.03.009

Izuma, K., & Adolphs, R. (2013). Social Manipulation of Preference in the Human Brain. Neuron, 78(3), 563–573. 10.1016/j.neuron.2013.03.023

Jin, R., Xia, T., Gawronski, B., & Hu, X. (2023). Attitudinal Effects of Stimulus Co-occurrence and Stimulus Relations: Sleep Supports Propositional Learning Via Memory Consolidation. Social Psychological and Personality Science, 14(1), 51–59. 10.1177/19485506211067673

Kendal, R. L., Boogert, N. J., Rendell, L., Laland, K. N., Webster, M., & Jones, P. L. (2018). Social Learning Strategies: Bridge-Building between Fields. Trends in Cognitive Sciences, 22(7), 651–665. 10.1016/j.tics.2018.04.003

Klinzing, J. G., Niethard, N., & Born, J. (2019). Mechanisms of systems memory consolidation during sleep. Nature Neuroscience, 22(10), 1598–1610. 10.1038/s41593-019-0467-3

Klucharev, V., Hytönen, K., Rijpkema, M., Smidts, A., & Fernández, G. (2009). Reinforcement Learning Signal Predicts Social Conformity. Neuron, 61(1), 140–151. 10.1016/j.neuron.2008.11.027

Kurdziel, L., Duclos, K., & Spencer, R. M. C. (2013). Sleep spindles in midday naps enhance learning in preschool children. Proceedings of the National Academy of Sciences, 110(43), 17267–17272. 10.1073/pnas.1306418110

Lehmann, M., Schreiner, T., Seifritz, E., & Rasch, B. (2016). Emotional arousal modulates oscillatory correlates of targeted memory reactivation during NREM, but not REM sleep. Scientific Reports, 6(1), 39229. 10.1038/srep39229

Liu, J., Xia, T., Chen, D., Yao, Z., Zhu, M., Antony, J. W., Lee, T. M. C., & Hu, X. (2023). Item-specific neural representations during human sleep support long-term memory. PLOS Biology, 21(11), e3002399. 10.1371/journal.pbio.3002399

Mednick, S. C., McDevitt, E. A., Walsh, J. K., Wamsley, E., Paulus, M., Kanady, J. C., & Drummond, S. P. A. (2013). The Critical Role of Sleep Spindles in Hippocampal-Dependent Memory: A Pharmacology Study. Journal of Neuroscience, 33(10), 4494– 4504. 10.1523/JNEUROSCI.3127-12.2013

Molho, C., Tybur, J. M., Van Lange, P. A. M., & Balliet, D. (2020). Direct and indirect punishment of norm violations in daily life. Nature Communications, 11(1), 3432. 10.1038/s41467-020-17286-2

Murty, V. P., FeldmanHall, O., Hunter, L. E., Phelps, E. A., & Davachi, L. (2016). Episodic memories predict adaptive value-based decision-making. Journal of Experimental Psychology: General, 145(5), 548–558. 10.1037/xge0000158

Nook, E. C., & Zaki, J. (2015). Social Norms Shift Behavioral and Neural Responses to Foods. Journal of Cognitive Neuroscience, 27(7), 1412–1426. 10.1162/jocn_a_00795

Oudiette, D., Antony, J. W., Creery, J. D., & Paller, K. A. (2013). The Role of Memory Reactivation during Wakefulness and Sleep in Determining Which Memories Endure. The Journal of Neuroscience, 33(15), 6672–6678. 10.1523/JNEUROSCI.5497-12.2013

Oudiette, D., & Paller, K. A. (2013). Upgrading the sleeping brain with targeted memory reactivation. Trends in Cognitive Sciences, 17(3), 142–149. 10.1016/j.tics.2013.01.006

Paller, K. A., Creery, J. D., & Schechtman, E. (2021). Memory and Sleep: How Sleep Cognition Can Change the Waking Mind for the Better. Annual Review of Psychology, 72(1), 123–150. 10.1146/annurev-psych-010419-050815

Pan, Y., Novembre, G., & Olsson, A. (2022). The Interpersonal Neuroscience of Social Learning. Perspectives on Psychological Science, 17(3), 680–695. 10.1177/17456916211008429

Paulhus, D. L. (1984). Two-component models of socially desirable responding. Journal of Personality and Social Psychology, 46(3), 598.

Payne, K., & Lundberg, K. (2014). The Affect Misattribution Procedure: Ten Years of Evidence on Reliability, Validity, and Mechanisms: Affect Misattribution Procedure. Social and Personality Psychology Compass, 8(12), 672–686. 10.1111/spc3.12148

Peirce, J., Gray, J. R., Simpson, S., MacAskill, M., Höchenberger, R., Sogo, H., Kastman, E., & Lindeløv, J. K. (2019). PsychoPy2: Experiments in behavior made easy. Behavior Research Methods, 51(1), 195–203.

Petzka, M., Chatburn, A., Charest, I., Balanos, G. M., & Staresina, B. P. (2022). Sleep spindles track cortical learning patterns for memory consolidation. Current Biology, 32(11), 2349–2356.e4. 10.1016/j.cub.2022.04.045

Plassmann, H., O’Doherty, J., & Rangel, A. (2007). Orbitofrontal Cortex Encodes Willingness to Pay in Everyday Economic Transactions. Journal of Neuroscience, 27(37), 9984–9988. 10.1523/JNEUROSCI.2131-07.2007

Rakowska, M., Abdellahi, M. E. A., Bagrowska, P., Navarrete, M., & Lewis, P. A. (2021). Long term effects of cueing procedural memory reactivation during NREM sleep. NeuroImage, 244, 118573. 10.1016/j.neuroimage.2021.118573

Rasch, B., & Born, J. (2013). About Sleep’s Role in Memory. Physiological Reviews, 93(2), 681–766. 10.1152/physrev.00032.2012

Raudenbush, B., & Frank, R. A. (1999). Assessing Food Neophobia: The Role of Stimulus Familiarity. Appetite, 32(2), 261–271. 10.1006/appe.1999.0229

Rihm, J. S., Diekelmann, S., Born, J., & Rasch, B. (2014). Reactivating Memories during Sleep by Odors: Odor Specificity and Associated Changes in Sleep Oscillations. Journal of Cognitive Neuroscience, 26(8), 1806–1818. 10.1162/jocn_a_00579

Rudoy, J. D., Voss, J. L., Westerberg, C. E., & Paller, K. A. (2009). Strengthening Individual Memories by Reactivating Them During Sleep. Science, 326(5956), 1079–1079. 10.1126/science.1179013

Schechtman, E., Antony, J. W., Lampe, A., Wilson, B. J., Norman, K. A., & Paller, K. A. (2021). Multiple memories can be simultaneously reactivated during sleep as effectively as a single memory. Communications Biology, 4(1), 1–13. 10.1038/s42003-020-01512-0

Schechtman, E., Heilberg, J., & Paller, K. A. (2023). Memory consolidation during sleep involves context reinstatement in humans. Cell Reports, 42(4), 112331. 10.1016/j.celrep.2023.112331

Schönauer, M., Alizadeh, S., Jamalabadi, H., Abraham, A., Pawlizki, A., & Gais, S. (2017). Decoding material-specific memory reprocessing during sleep in humans. Nature Communications, 8(1), 15404. 10.1038/ncomms15404

Schreiner, T., Lehmann, M., & Rasch, B. (2015). Auditory feedback blocks memory benefits of cueing during sleep. Nature Communications, 6(1), 8729. 10.1038/ncomms9729

Schreiner, T., Petzka, M., Staudigl, T., & Staresina, B. P. (2021). Endogenous memory reactivation during sleep in humans is clocked by slow oscillation-spindle complexes. Nature Communications, 12(1), 3112. 10.1038/s41467-021-23520-2

Shanahan, L. K., Gjorgieva, E., Paller, K. A., Kahnt, T., & Gottfried, J. A. (2018). Odor-evoked category reactivation in human ventromedial prefrontal cortex during sleep promotes memory consolidation. eLife, 7, e39681. 10.7554/eLife.39681

Sterpenich, V., Van Schie, M. K. M., Catsiyannis, M., Ramyead, A., Perrig, S., Yang, H.-D., Van De Ville, D., & Schwartz, S. (2021). Reward biases spontaneous neural reactivation during sleep. Nature Communications, 12(1), 4162. 10.1038/s41467-021-24357-5

Strachan, J. W. A., Guttesen, A. á V., Smith, A. K., Gaskell, M. G., Tipper, S. P., & Cairney, S. A. (2020). Investigating the formation and consolidation of incidentally learned trust. *Journal of Experimental Psychology: Learning*, Memory, and Cognition, 46(4), 684–698. 10.1037/xlm0000752

Templeton, E. M., Stanton, M. V., & Zaki, J. (2016). Social Norms Shift Preferences for Healthy and Unhealthy Foods. PLOS ONE, 11(11), e0166286. 10.1371/journal.pone.0166286

Vallat, R., & Walker, M. P. (2021). An open-source, high-performance tool for automated sleep staging. eLife, 10, e70092. 10.7554/eLife.70092

Venkatraman, V., Huettel, S. A., Chuah, L. Y. M., Payne, J. W., & Chee, M. W. L. (2011). Sleep Deprivation Biases the Neural Mechanisms Underlying Economic Preferences. Journal of Neuroscience, 31(10), 3712–3718. 10.1523/JNEUROSCI.4407-10.2011

Wilhelm, I., Diekelmann, S., Molzow, I., Ayoub, A., Molle, M., & Born, J. (2011). Sleep Selectively Enhances Memory Expected to Be of Future Relevance. Journal of Neuroscience, 31(5), 1563–1569. 10.1523/JNEUROSCI.3575-10.2011

Wimmer, G. E., & Büchel, C. (2016). Reactivation of Reward-Related Patterns from Single Past Episodes Supports Memory-Based Decision Making. Journal of Neuroscience, 36(10), 2868–2880. 10.1523/JNEUROSCI.3433-15.2016

Wimmer, G. E., & Shohamy, D. (2012). Preference by Association: How Memory Mechanisms in the Hippocampus Bias Decisions. Science, 338(6104), 270–273. 10.1126/science.1223252

Xia, T., Antony, J. W., Paller, K. A., & Hu, X. (2023). Targeted memory reactivation during sleep influences social bias as a function of slow-oscillation phase and delta power. Psychophysiology, 60(5), e14224. 10.1111/psyp.14224

Xia, T., Chen, D., Zeng, S., Yao, Z., Liu, J., Qin, S., Paller, K. A., Torres-Platas, S. G., Antony, J. W., & Hu, X. (2023). Aversive memories can be weakened during human sleep via the reactivation of positive interfering memories. bioRxiv, doi: 10.1101/2023.12.05.570072

Xia, T., Yao, Z., Guo, X., Liu, J., Chen, D., Liu, Q., Paller, K. A., & Hu, X. (2023). Updating memories of unwanted emotions during human sleep. Current Biology, 33(2), 309–320.e5. 10.1016/j.cub.2022.12.004

Yu, H., Siegel, J. Z., Clithero, J. A., & Crockett, M. J. (2021). How peer influence shapes value computation in moral decision-making. Cognition, 211, 104641. 10.1016/j.cognition.2021.104641

Yuksel, C., Denis, D., Coleman, J., Oh, A., Cox, R., Morgan, A., Sato, E., & Stickgold, R. (2023). Emotional memories are enhanced when reactivated in slow wave sleep, but impaired when reactivated in REM. bioRxiv. doi: 10.1101/2023.03.01.530661

Zaki, J., Schirmer, J., & Mitchell, J. P. (2011). Social Influence Modulates the Neural Computation of Value. Psychological Science, 22(7), 894–900. 10.1177/0956797611411057

Zhang, L., & Gläscher, J. (2020). A brain network supporting social influences in human decision-making. Science Advances, 6(34), eabb4159. 10.1126/sciadv.abb4159

Züst, M. A., Ruch, S., Wiest, R., & Henke, K. (2019). Implicit Vocabulary Learning during Sleep Is Bound to Slow-Wave Peaks. Current Biology, 29(4), 541–553.e7. 10.1016/j.cub.2018.12.038

