## Supplementary material for "Modulating social learning-induced evaluation updating during human sleep": Figure S1; Figure S2; Figure S3; Table S1; Table S2; Table S3

**This file includes:**

Figure S1 to S3

Table S1 to S3

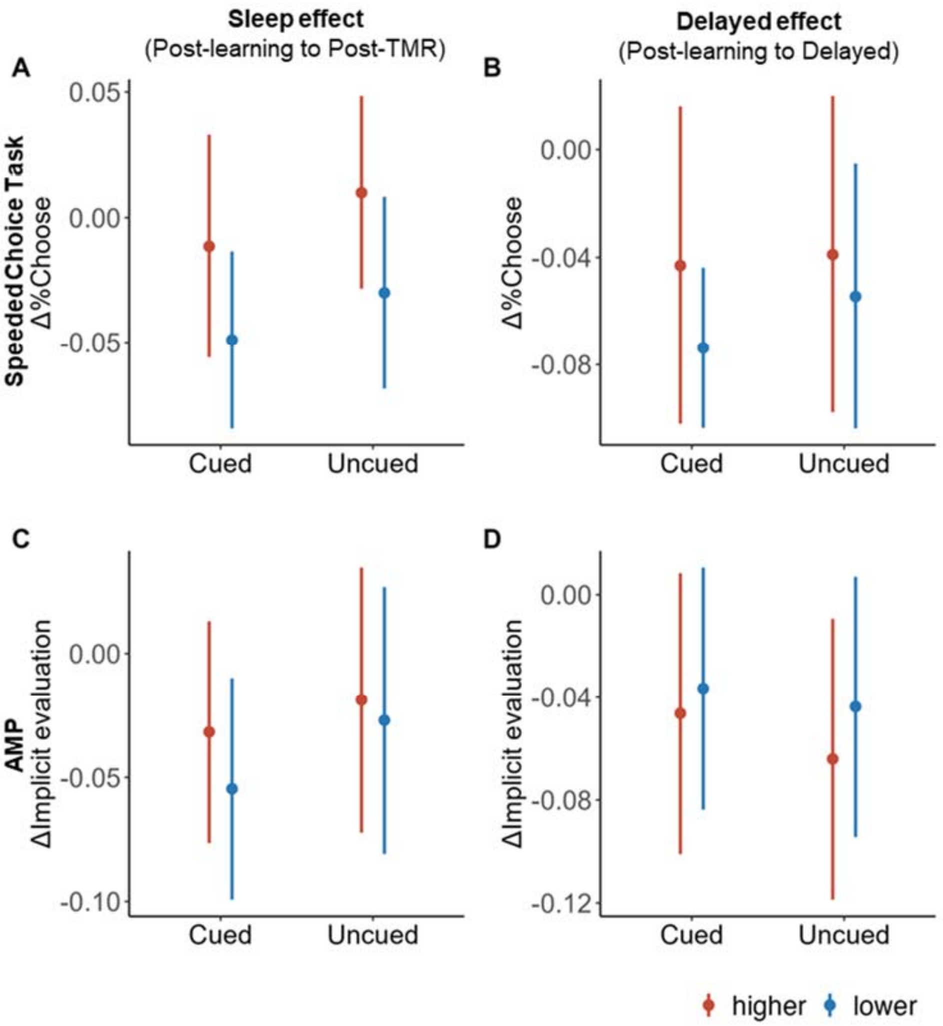


**Figure S1:** Impact of TMR and social learning on Δ%Choose in the speeded choice (A) from post-learning to post-TMR phase, and (B) from post-learning to delayed phase. Impact of TMR and social learning on ΔImplicit evaluation in the AMP task (A) from post-learning to post-TMR phase, and (B) from post-learning to delayed phase.

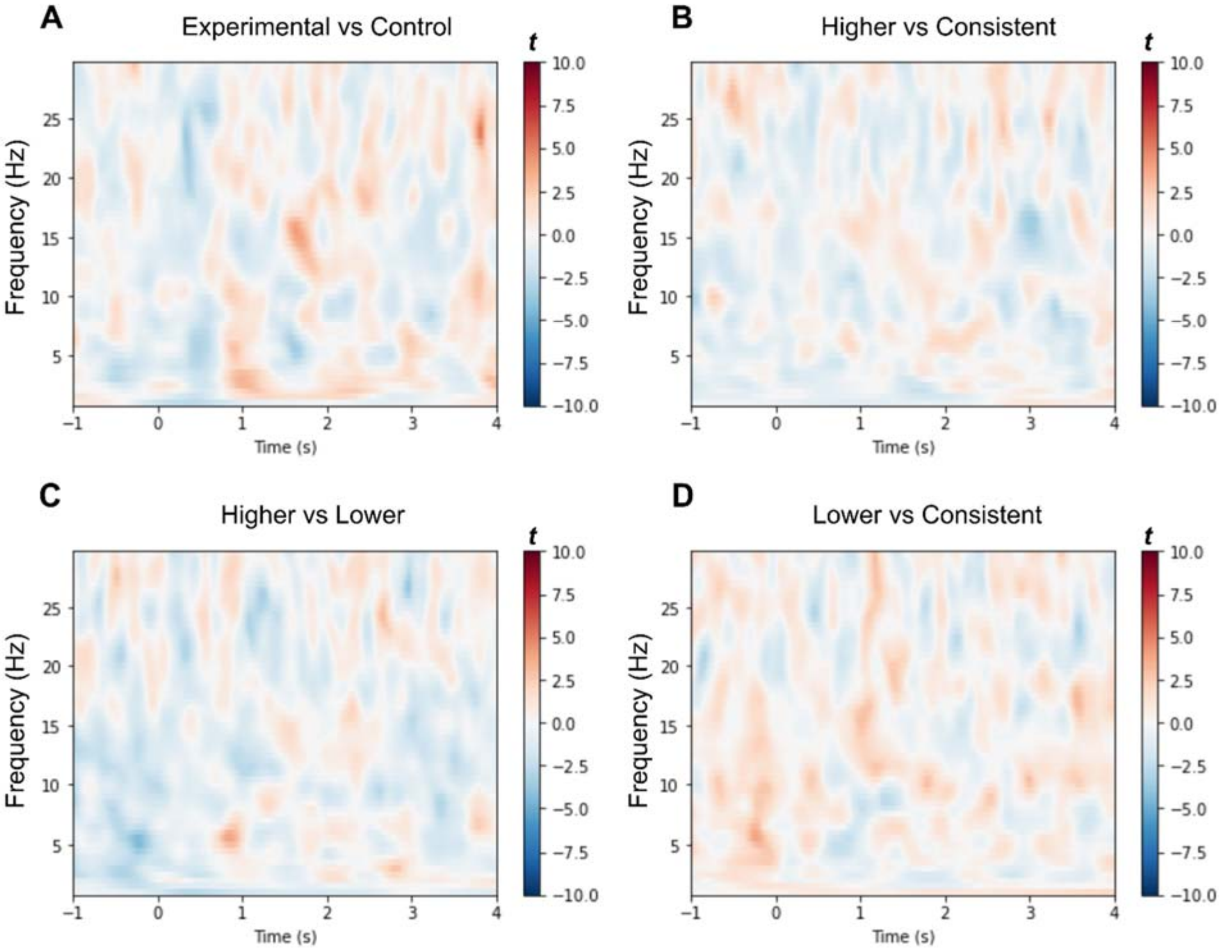


**Figure S2:** Cue-elicited EEG power comparing A) experimental cue vs. control cues, B) “Higher” cue vs. “Consistent” cue, C) “Higher” cue vs. “Lower” cue, and D) “Lower” cue vs. “Consistent” cue. No significant cluster was found across all four comparisons ($p_{cluster}$s > .217).

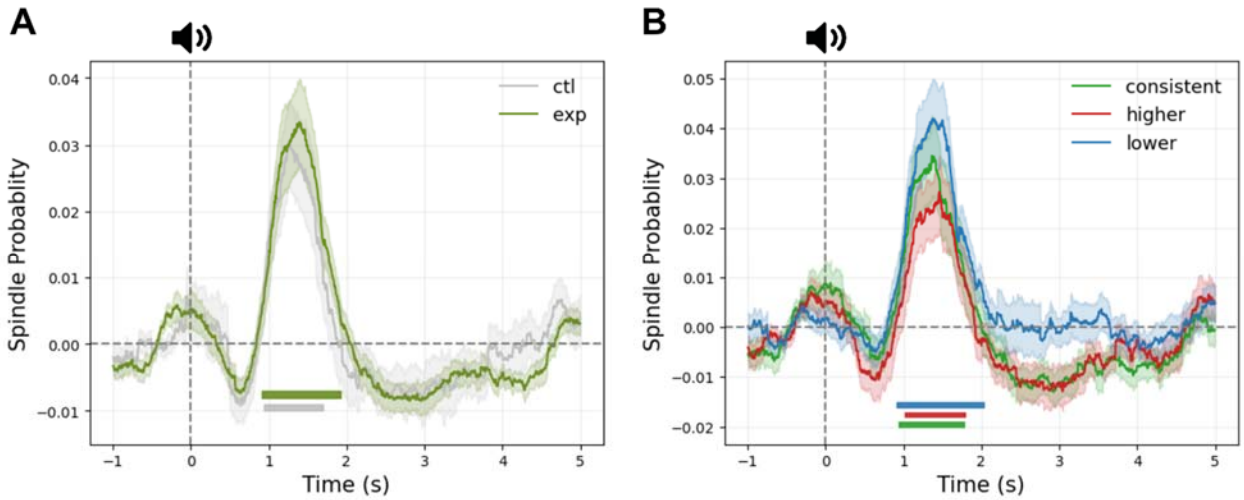


**Figure S3:** Spindle Probability (A) between Control and Experimental conditions, and (B) among Higher, Lower, and Consistent conditions. The shaded part indicated standard errors. The colored line indicated the significant clusters when comparing the spindle probability with the baseline probability.

Table S1. Baseline preference and familiarity (Mean [S.D.]) across conditions.

| Measurements | Preference | Familiarity |
|---|---|---|
| *Cued* | | |
| Higher | 6.05 (1.02) | 5.56 (1.25) |
| Consistent | 6.07 (1.03) | 5.59 (1.25) |
| Lower | 6.09 (0.98) | 5.90 (1.32) |
| *Uncued* | | |
| Higher | 6.05 (1.02) | 5.67 (1.08) |
| Consistent | 6.13 (1.06) | 5.74 (1.17) |
| Lower | 6.09 (1.04) | 5.84 (1.08) |
| *Statistics* | | |
| Main effect of TMR | $F$ (1, 33) = 0.86, p = .361 | $F$ (1, 33) = 0.22, $p$ = .644 |
| Main effect of feedback | $F$ (1.90, 62.72) = 2.58, p = .087 | $F$ (1.83, 60.24) = 1.89, $p$ = .163 |
| Interaction effect | $F$ (1.86, 61.52) = 0.87, p = .419 | $F$ (1.78, 58.90) = 0.30, $p$ = .715 |

Table S2. Remained TMR epoch numbers.

| Condition | Mean | S.D. |
|---|---|---|
| Higher | 113.82 | 31.21 |
| Lower | 113.91 | 31.26 |
| Consistent | 113.97 | 30.84 |
| Control | 114.18 | 30.94 |
| In total | 455.88 | 124.25 |

Table S3. Sleep Measurements.

| Measurement | Mean | S.D. |
|---|---|---|
| TIB | 483.24 | 25.60 |
| SPT | 467.57 | 32.12 |
| WASO | 27.72 | 25.14 |
| TST | 439.85 | 41.81 |
| N1 | 20.76 | 9.41 |
| N2 | 215.97 | 26.29 |
| N3 | 100.75 | 26.90 |
| REM | 102.36 | 24.39 |
| NREM | 337.49 | 31.61 |
| SOL | 14.21 | 17.16 |
| Lat_N1 | 22.94 | 52.63 |
| Lat_N2 | 19.32 | 17.80 |
| Lat_N3 | 28.36 | 19.71 |
| Lat_REM | 92.39 | 30.96 |
| %N1 | 4.80 | 2.28 |
| %N2 | 49.32 | 6.12 |
| %N3 | 22.76 | 5.21 |
| %REM | 23.12 | 4.64 |
| %NREM | 76.88 | 4.64 |
| SE | 91.00 | 6.81 |
| SME | 94.03 | 5.38 |
| Stability | 0.92 | 0.02 |

*Note*: We provided the interaction effect of linear-mixed model analyses using sleep measurements to predict the ΔPreference of cued items from post-learning to post-TMR phases. TIB = Time in Bed. SPT = Sleep Period Time. WASO = Wake After Sleep Onset. TST = Total Sleep Time. N1, N2, N3, and REM: Sleep stages duration. NREM = N1 + N2 + N3. SOL = Sleep Onset Latency. Lat_N1, N2, N3, REM: latencies of sleep stages from the beginning of the record. %(N1, N2, N3, REM): Sleep stages duration expressed in percentages of TST. S.E. = Sleep Efficiency. SME: Sleep Maintenance Efficiency. Stability = diagonal value in the transition matrix.
